# A human stem cell resource to decipher the biochemical and cellular basis of neurodevelopmental defects in Lowe Syndrome

**DOI:** 10.1101/2021.08.19.456986

**Authors:** Bilal M. Akhtar, Priyanka Bhatia, Shubhra Acharya, Sanjeev Sharma, Yojet Sharma, BS Aswathy, Kavina Ganapathy, Anil Vasudevan, Padinjat Raghu

## Abstract

Human brain development is a complex process where multiple cellular and developmental events are co-ordinated to generate normal structure and function. Alteration in any of these events can impact brain development, manifesting clinically as neurodevelopmental disorders. Human genetic disorders of lipid metabolism often present with features of altered brain function. Lowe syndrome (LS), is a X-linked recessive disease with features of altered brain function. LS results from mutations in *OCRL1* that encodes a phosphoinositide 5-phosphatase enzyme. However, the cellular mechanisms by which loss of *OCRL1* leads to brain defects remain unknown. Human brain development involves several cellular and developmental features not conserved in other species and understanding such mechanisms remains a challenge. Rodent models of LS have been generated, but failed to recapitulate features of the human disease. Here we describe the generation of human stem cell lines from LS patients. Further, we present biochemical characterization of lipid metabolism in patient cell lines and demonstrate their use as a “*disease-in-a-dish*” model for understanding the mechanism by which loss of *OCRL1* leads to altered cellular and physiological brain development.

## Introduction

Phosphoinositides are key regulators of the organization and function of eukaryotic cells (Schink et al., 2016). Of the seven species of phosphoinositides, phosphatidylinositol 4,5-bisphosphate [PI(4,5)P_2_] is the most abundant and is required to regulate multiple sub-cellular processes including membrane turnover, cytoskeletal function and the organization of the plasma membrane (Kolay et al., 2016). PI(4,5)P_2_ exerts its control over cellular functions both through binding and allosteric regulation of protein activity and also through its ability to serve as a substrate for phospholipase C (PLC) and Class I PI3K signalling (Katan and Cockcroft, 2020). Therefore, the accurate regulation of PI(4,5)P_2_ levels at cellular membranes is critical for normal function. PI(4,5)P_2_ levels are regulated through enzymes that regulate its synthesis, the phosphatidylinositol 4-phosphate 5-kinases (PIP5K) (van den Bout and Divecha, 2009) and also by enzymes that catalyze its metabolism. In addition to PLC and Class I PI3K that utilise PI(4,5)P_2_ to generate signalling molecules, lipid phosphatases that can dephosphorylate PI(4,5)P_2_ have also been described. These include 4-phosphatase enzymes that generate PI5P, but their function *in vivo* remains unclear (Ungewickell et al., 2005). A large family of 5-phosphatases that can dephosphorylate PI(4,5)P_2_ at position 5 to generate phosphatidylinositol 4-phosphate (PI4P) have been described (Ooms et al., 2009). This 5-phosphatase activity is encoded in all major eukaryotic genomes, with mammalian genomes encoding upto ten genes for this family of proteins; many of these gene-products have been linked to human diseases (Ramos et al., 2019). The significance of encoding a single enzyme activity through such a large gene family remains to be understood.

The Oculocerebrorenal syndrome of Lowe gene (*OCRL*) gene encodes a 901 amino acid inositol polyphosphate 5-phosphatase enzyme, that is able to catalyse the removal of the 5′ phosphate from PI(4,5)P_2_ to generate PI4P. *OCRL* was originally identified as the gene underlying the human inherited disease Lowe Syndrome (LS) (Attree et al., 1992). In human patients with LS, sequencing studies have revealed a large diversity of mutations in *OCRL*, including deleterious missense and nonsense mutations in all of the major domains of the protein including the 5-phosphatase domain, PH, ASH and RhoGAP domain (Staiano et al., 2015). *OCRL* is widely expressed across many human tissues or organs and at all stages of life (Raghu et al., 2019). The function of OCRL has been studied by overexpression or depletion in a number of common human cell lines and the protein has been reported to localize to and affect the function of many cellular organelles (Mehta et al., 2014).

LS is a rare (∼ 1/500,000 males), X-linked, recessive disorder characterized by the triad of congenital cataracts, intellectual or neurodevelopmental impairment and proximal renal tubular dysfunction (https://omim.org/entry/300535) (Bökenkamp and Ludwig, 2016; De Matteis et al., 2017). The brain phenotypes in LS include delayed and impaired cognitive milestones, hypotonia, febrile seizures and hyperechoic changes in the periventricular zone of the cerebral cortex. There are two enigmatic and unresolved observations in relation to the clinical presentation of LS: (i) Although LS is a monogenic disorder, there is substantial variability in the clinical presentation between individual patients, even in those individuals with mutations in *OCRL* with equivalent molecular consequences (e.g truncating nonsense mutations prior to the start of the phosphatase domain). For e.g, while some patients present with severe neurodevelopmental phenotypes, others show relatively mild deficits in brain function. These disparities suggest that in addition to the mutation in *OCRL*, other changes in the genetic background of the individual may impact the clinical outcome of loss of *OCRL*. (ii) Although *OCRL* is widely expressed in human tissues, it remains a mystery as to why only three organs are affected in LS, namely the eye, brain and kidney. One possibility is that the phenotypic changes seen in human patients may arise due to the requirement of OCRL only in specific cell types of the affected organs. Therefore, the relevant cellular changes may only be seen when studying eye, brain or renal tissue during development. Although rodent models of *OCRL* were generated, they failed to show phenotypes that recapitulate the human disease (Jänne et al., 1998). A zebrafish model of *OCRL* depletion has been generated that recapitulates some aspects of the human phenotype but there remains a lack of models that allow the brain phenotype to be studied (Ramirez et al., 2012). A limited number of studies have been done on LS fibroblasts and renal biopsies, but there is presently no understanding of how loss of *OCRL* leads to neurodevelopmental phenotypes. Thus, there is a requirement for a model system in which the cellular and physiological changes in the brain during development can be studied.

One possible route to obtaining a suitable model system arises from the ability to use modern stem cell technology to generate human induced pluripotent stem cells (hiPSC) (Shi et al., 2017) from the somatic tissues of patients with LS. These hiPSC can then be differentiated into specific adult tissues and the trajectory of development along with the cellular and molecular changes in any particular patient derived line can be analysed (Russo et al., 2015; Zeng et al., 2014). In the context of LS, a limited number of studies have reported individual hiPSC lines derived from patients (Barnes et al., 2018; Hsieh et al., 2018; Liu et al., 2021; Qian et al., 2021). In this study, we present the generation of hiPSC lines and neural derivatives from a family with a unique genetic structure and clinical features that should allow an understanding of the cellular basis of the neurodevelopmental phenotype in LS. We also present a biochemical analysis of phosphoinositide levels and an insight into the biochemical compensation for the loss of the 5-phosphatase activity of OCRL in LS cells.

## Results

### Generation of hiPSCs from a family with Lowe syndrome

For this study, we selected a family whose genetic structure (Fig. 1A i) is uniquely suited for the analysis of neurodevelopmental phenotypes in LS (Ahmed P et al., 2021). Briefly, in this family, the patients with LS are children of two female siblings, both of whom carry the identical mutation in *OCRL*. While all three children show the triad of eye, renal and brain phenotypes characteristic of LS, the brain phenotype of LSPH004 is much more severe than that of the identical twins LSPH002 and LSPH003. To understand the cellular and developmental mechanisms that underlie this neurodevelopmental defect, we generated hiPSC from each of these patients. Peripheral blood mononuclear cells (PBMC) were isolated from each patient, immortalized into lymphoblastoid cells lines (LCLs) which were then reprogrammed to generate hiPSC (Iyer et al., 2018) (Fig. 1A ii). The hiPSC lines so generated did not express the OCRL protein as determined by immunocytochemistry with an antibody to OCRL (Fig. 1B). The hiPSC lines showed expression of pluripotency markers SOX2 and SSEA4 as determined by immunocytochemistry (Fig. 1C) and SSEA4 and OCT4 as determined by single cell quantitative fluorescence activated cell sorting (FACS) analysis (Supplementary Fig. 1A-C). We differentiated these hiPSC into embryoid bodies and established their ability to differentiate into each of the three germ layers by detecting the expression of transcripts characteristic of each layer (SOX1, Nestin-ectoderm, Nodal-mesoderm and GATA4-endoderm) (Fig. 1D). All hiPSC lines were determined to be of a normal karyotype (Fig. 1E, Supplementary Fig. 2D, E) and short tandem repeat analysis was used to determine and track the identity of each cell line (Supplementary Table 1). These hiPSC lines offer a unique resource from which tissue specific differentiation, for example into brain tissue can be carried out; a comparison of the cellular and molecular differences between control and patient derived hiPSC lines during brain development can provide important insights into how loss of OCRL results in altered neurodevelopment.

**Figure 1.**
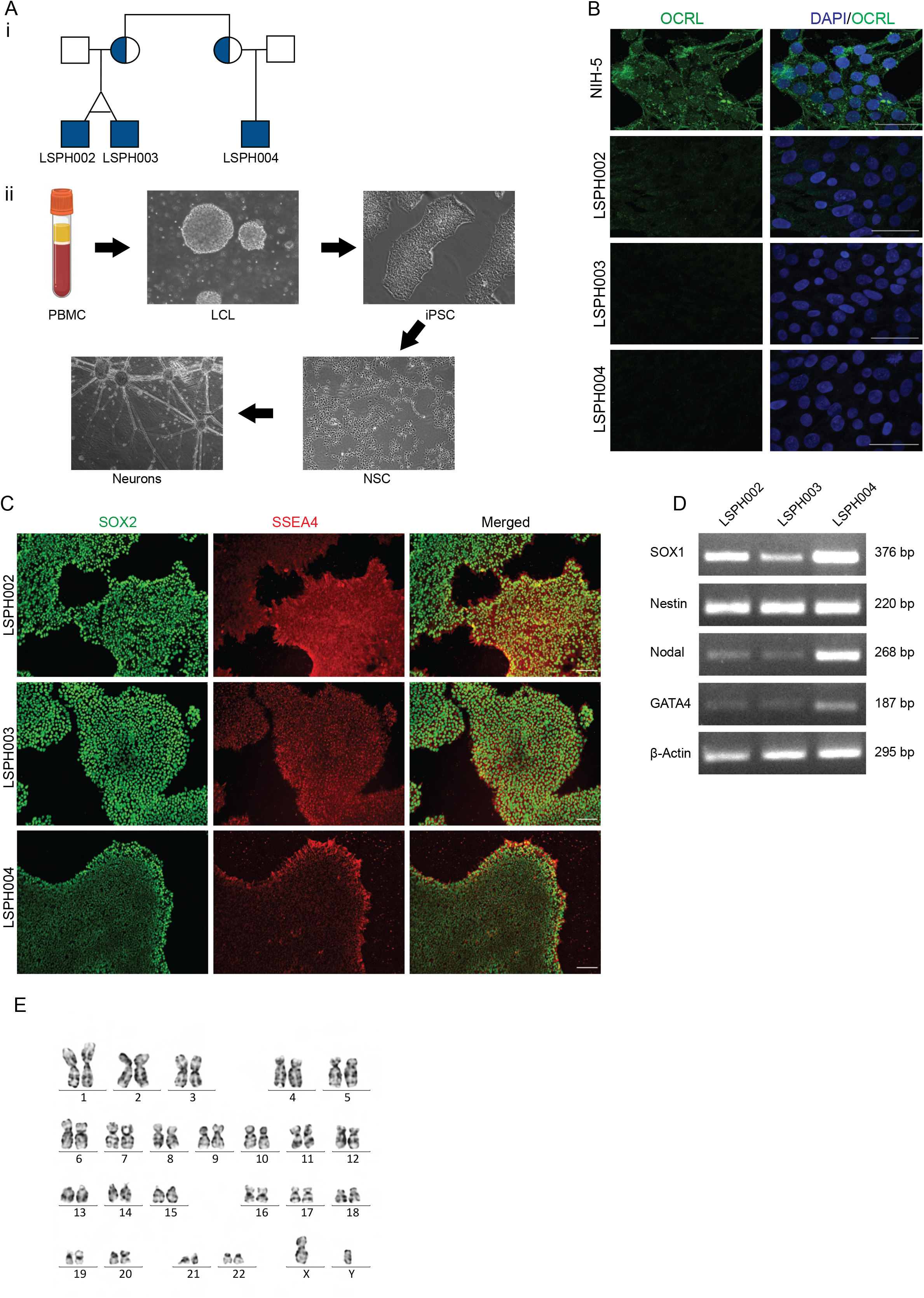
Generation and characterization of hiPSCs derived from Lowe syndrome patients. (A) i)Pedigree structure of the studied family. Shaded circles indicate heterozygous carrier mothers and shaded squares indicate hemizygous patients. ii) Steps in generation of iPSC, NSC and Neurons from LCL. (B) Immunocytochemistry of hiPSC colonies showing expression of OCRL (green) in control line NIH5 and absent in the patient lines (LSPH002, LSPH003, LSPH004). Nuclear stain DAPI (blue) Scale bar: 50μm. (C) Immunocytochemistry of hiPSC colonies showing presence of pluripotency nuclear marker SOX2 (green) and of pluripotency surface marker SSEA4 (red). Scale bar: 200μm. (D) Expression of three germ lineage transcripts SOX1 and Nestin (Ectoderm), Nodal (Mesoderm), GATA4 (Endoderm), β-actin as control by RT-PCR from embryoid bodies derived from LSPH002, LSPH003, LSPH004 hiPSC. (E) Karyogram depicting normal karyotype 46(X,Y) of the hiPSC line LSPH004

**Figure 2.**
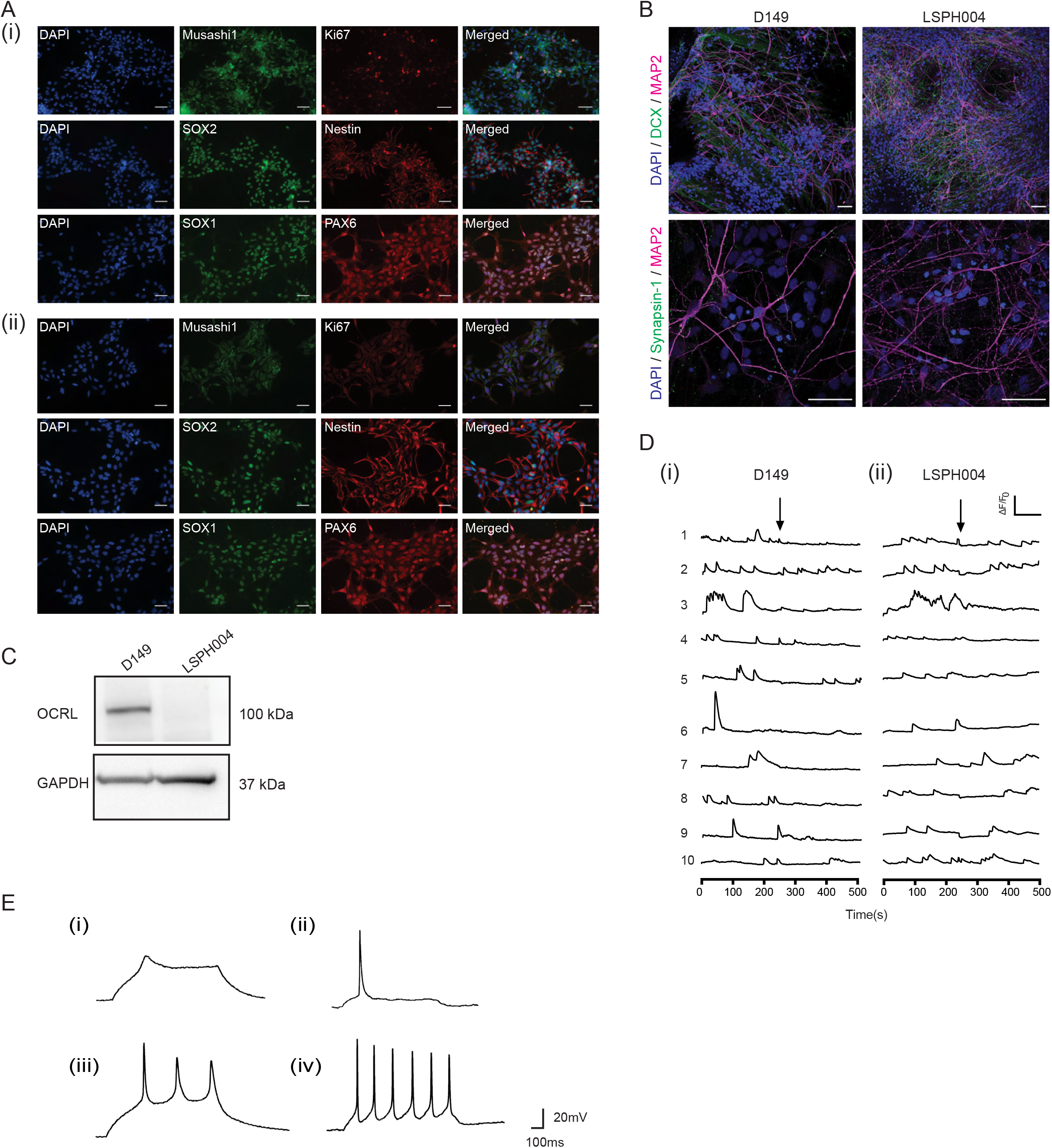
Characterization of Neural Stem Cells and Neurons derived from Lowe syndrome hiPSC. (A) Immunocytochemistry of Neural Stem Cells (i) D149 (control) (ii) LSPH004 (patient) showing the expression of the NSC markers SOX1, SOX2, PAX-6, Nestin, Mushashi-1 and the proliferation marker Ki-67. Nucleus stained with DAPI. Scale bar: 50μm (B) Immunofluorescence images (maximum intensity projections) of D149 and LSPH004 neurons at 30 DIV differentiated from respective NSCs. The cells were stained with the following neuronal markers: (i, ii) DCX (*green*, immature neuronal marker) and MAP2 (*magenta*, mature neuronal marker); (iii, iv) synapsin-1 (*green*) and MAP2 (*magenta*); followed by counterstaining with DAPI (*blue*). Scale bar: 50μm. (C) Western blot showing expression of OCRL protein in lysates from 30 DIV Neurons in the control line D149 and its absence in patient line LSPH004. GAPDH was used as a loading control. (D) Calcium transients recorded from 30 DIV D149 (i) and LSPH004 (ii) neurons are shown. Each panel shows [Ca^2+^]_i_ traces from individual cells in the dish. Y-axis shows normalized fluorescence intensity ΔF/F_0_ and X-axis is time in seconds. The baseline recording for 4 mins, followed by addition of 10μM tetrodotoxin (TTX) (as indicated by the arrows). (E) Evoked action potentials (AP) in neurons differentiated from NSC recorded using whole-cell patch clamp electrophysiology at 10, 20, 30 and 40 DIV. The characteristic feature of action potentials recorded from control D149 cells at each time point is shown. (i) immature AP at 10 DIV, single action potential at 20 DIV (ii) and (iii) multiple AP on 30 DIV (iv) Most neurons exhibited repetitive firing by 40 DIV.

### Generation of Neural Stem Cells (NSC) from hiPSC

During the development of the brain, a key step in the conversion of pluripotent, early embryonic stem cells into brain cells is the formation of neural stem cells (NSC) which then both divide and differentiate to generate the different cell types of the brain. Thus, with the exception of microglia, NSC can be differentiated into all cell types in the brain. Since the neurodevelopmental phenotype was strongest in LSPH004, we generated NSC to understand the brain development phenotype of LS. As a control, we generated NSC from D149 hiPSC (Iyer et al., 2018) that was originally derived from an unaffected individual of similar population background (Fig 1A ii). The NSC so generated were characterized by confirming expression of established NSC protein markers such as Nestin, SOX1, SOX2, PAX6 and Musashi-1 along with the proliferation marker Ki-67 (Fig. 2A). They were also confirmed to be karyotypically normal (Supplementary Fig. 3A-B). Lastly, these NSC were differentiated into cultures of forebrain cortical neurons. The neurons so generated showed the characteristic morphology and molecular markers of neuronal development including MAP2, DCX and Synapsin-1 (Fig. 2B) as previously reported (Sharma et al., 2020). Western blot analysis of protein extracts from these lysates revealed the band corresponding to OCRL in wild type neurons and absent in LSPH004 (Fig. 2C). To test the physiological status of these neurons, we monitored them for the presence of intracellular calcium transients [Ca^2+^]_i_, a characteristic feature associated with neuronal development (Rosenberg and Spitzer, 2011). We found that in 30 days *in vitro* (DIV) cultures, robust [Ca^2+^]_i_ transients were observed in neurons derived from both D149 (Fig. 2D i) and LSPH004 (Fig. 2D ii). In addition, we also monitored the development of electrical activity in the differentiating neurons as a function of age *in vitro* using whole cell, patch clamp electrophysiology in both D149 and LSPH004. For example, in 10 DIV cultures of D149, abortive action potentials were noted (Fig. 2E i); by 20 DIV single action potentials were seen (Fig. 2E ii), by 30 DIV multiple action potentials were noted (Fig. 2E iii) and by 40 DIV repetitive firing was observed (Fig. 2E iv).

**Figure 3.**
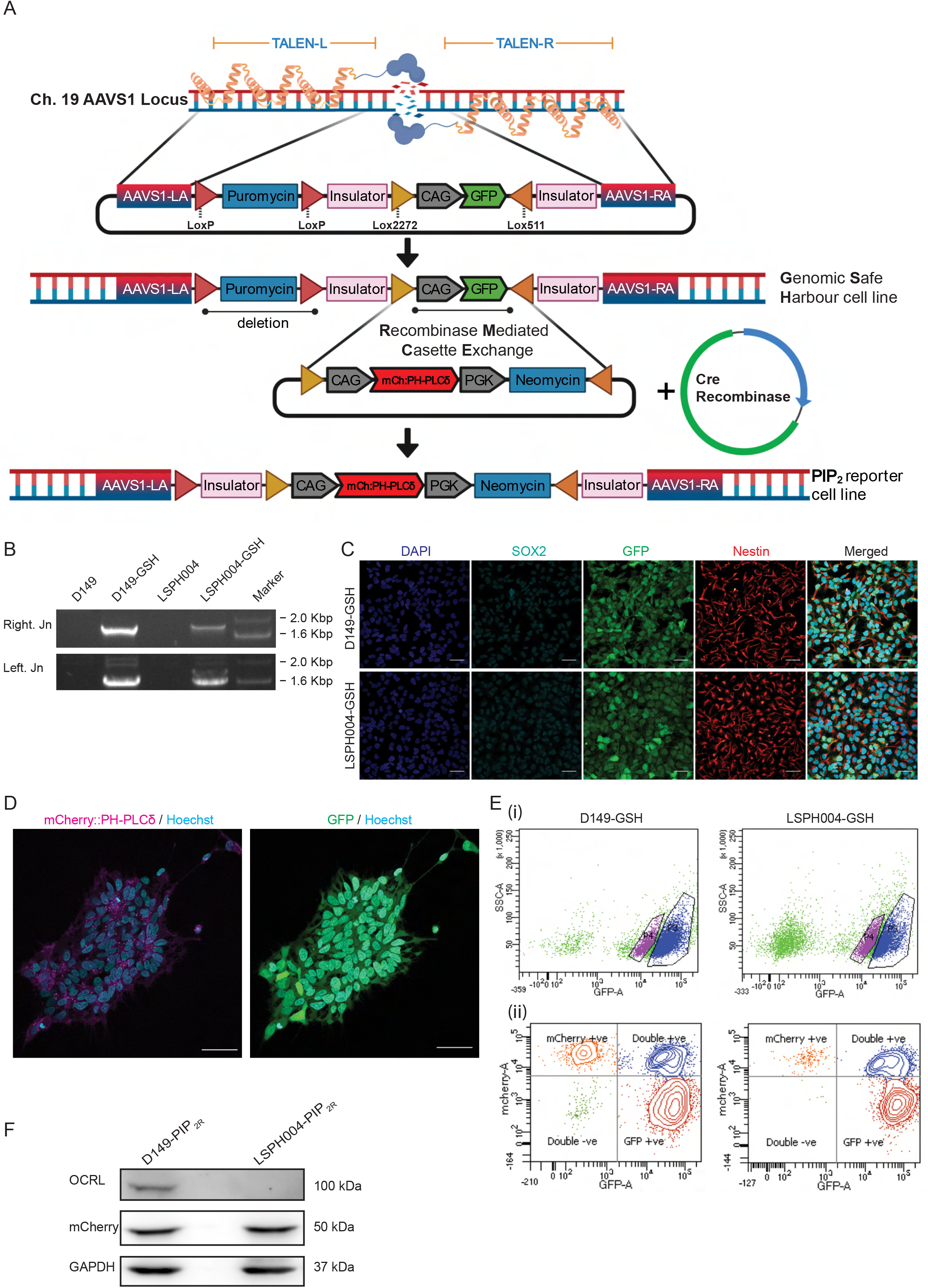
Generation of genomic safe harbour lines for stable expression of protein biosensors. (A) A schematic showing the mechanism for generating the reporter line using Genomic Safe Harbor-RMCE (Recombinase Mediated Cassette Exchange) approach. Abbreviations: TALEN-Transcription activator-like effector nucleases (Left and Right), AAVS1-Adeno-Associated Virus Integration Site 1 (LA-Left Arm, RA-Right Arm). (B) Validation of LoxP-CAG-GFP (ZYP037) insert into AAVS1 genomic locus using junction PCR. Amplicon of the expected size validating the right and left junction are shown. (C) Characterization of Genomic Safe Harbour lines D149-GSH and LSPH004-GSH: Expression of the NSC markers Nestin and SOX2 detected by Immunocytochemistry. GFP expression from the safe harbour construct marker is also shown. Scale bar: 40μm. (D) Representative confocal image (maximum z-projection) showing expression and localization of mCherry::PH-PLCδ (magenta) and cytosolic GFP (green) in D149 PIP_2R_ reporter line colonies. Nucleus stained using Hoechst (cyan). Scale bar: 50 μm. (E) Representative flow cytometry data to illustrate the gating strategy for FACS purification of i) GFP positive cells for Genomic Safe Harbour lines, to select for healthy GFP positive cells, side scatter (SSC-A, log, Y-axis) and signal from the excitation of cells with the 488-nm laser (GFP-A, log, X-axis) are plotted. ii) mCherry positive cells for PIP_2_ reporter lines were selected for by plotting signal from the excitation of cells with the 568-nm laser (mCherry-A, log, Y-axis) against signal from the excitation of cells with the 488-nm laser (GFP-A, log, X-axis). (F) Western blot showing OCRL (detected using the OCRL antibody), mCherry::PH-PLCδ (detected using the mCherry antibody) in Neural Stem Cell PIP_2R_ reporter lines. GAPDH was used as a loading control.

### Tools for controlled expression of proteins in NSC

In order to understand the cellular basis of the neurodevelopmental defect in LS, it is essential to monitor in real time, ongoing cellular and molecular processes using protein based reporters such as those used to monitor phosphoinositide turnover at specific cellular membranes (Hammond and Balla, 2015). Likewise, in order to understand the function of specific domains of OCRL including the 5- phosphatase domain, it will be necessary to achieve carefully controlled reconstitution of OCRL variants in LS cells during development. Current methods of protein expression in neural cells, such as lentiviral mediated transduction of transgenes, result in variable levels of expression between cell lines and experiments. One strategy for the controlled expression of proteins in stem cells is integration of genetic constructs at specific locations in the genome that have been found to be suitable for expression of a transgene/biosensor without any major adverse consequences due to insertion (Pei et al., 2015). Briefly, transcription activator-like effector nucleases (TALENs) are used to insert a Lox cassette at adeno-associated virus integration site 1 (AAVS1) [referred to as Genomic safe harbour sites (GSH)] locus on chromosome 19 thus generating a master cell line for subsequent insertion of transgenes at this location. Transgenes of interest, under suitable promoters, can be inserted into this safe harbour site by Lox- P/Cre-recombinase mediated cassette exchange (RMCE) (Fig. 3A). The advantage of this approach is that there is no rapid loss of transgene expression after transfection and no variation in copy number of transgenes between experiments or cell lines as might occur with transient transfection or lentiviral transduction of transgenes.

Using this approach, we inserted GSHs into D149 and LSPH004 NSCs. Insertion of GSHs was monitored by observing GFP expression and using a junction PCR (Fig. 3B) and the NSC line so generated continued to express characteristic protein markers (Fig. 3C). RMCE was used to insert a Lox cassette into GSHs of D149 (control) and LSPH004 (patient) NSCs; this cassette included the cDNA for a protein probe for PI(4,5)P_2_, the PH domain of PLCδ fused to mCherry (mCherry:: PH- PLCδ) (Várnai and Balla, 1998). In this cassette, the mCherry::PH-PLCδ is expressed under a CAG promoter that is active in NSC. NSC in which the mCherry::PH-PLCδ transgene is recombined into the GSH show mCherry fluorescence, NSC in which only one of two copies of the GSH have undergone RMCE show both mCherry and GFP fluorescence and cells in which no recombination has taken place show only GFP fluorescence (Fig. 3D). We purified NSC with mCherry fluorescence only using FACS (Fig. 3E). Western blot analysis on these purified cells showed that they express a protein with a M_r_ corresponding to that of mCherry::PH-PLCδ (Fig. 3F).

### Impact of OCRL depletion on total cellular PI(4,5)P_2_

Since OCRL is a PI(4,5)P_2_ 5-phosphatase, it is expected that cellular PI(4,5)P_2_ levels might be elevated and PI4P levels reduced in LS patient cells. To test this, we extracted total lipids from hiPSC and NSC of both control and patient derived LSPH004 cells. PIP_2_ levels were measured using liquid chromatography coupled with mass spectrometry (LCMS) from whole cell lysates (Sharma et al., 2019); although this method cannot distinguish between positional isomers of PIP_2_, the majority is expected to be PI(4,5)P_2_. In hiPSCs, a significant increase was seen in the total PIP (Fig. 4A) and PIP_2_ (Fig. 4B) mass in LSPH004 compared to control (Fig. 4A-B). In experiments with NSCs, we compared D149 with LSPH004 and found no difference in the total PIP mass (Fig. 4C) or PIP_2_ mass (Fig. 4D) between these two lines. Thus, loss of OCRL results in a modest change in PI4P and PI(4,5)P_2_ mass in LS patient derived hiPSC.

**Figure 4.**
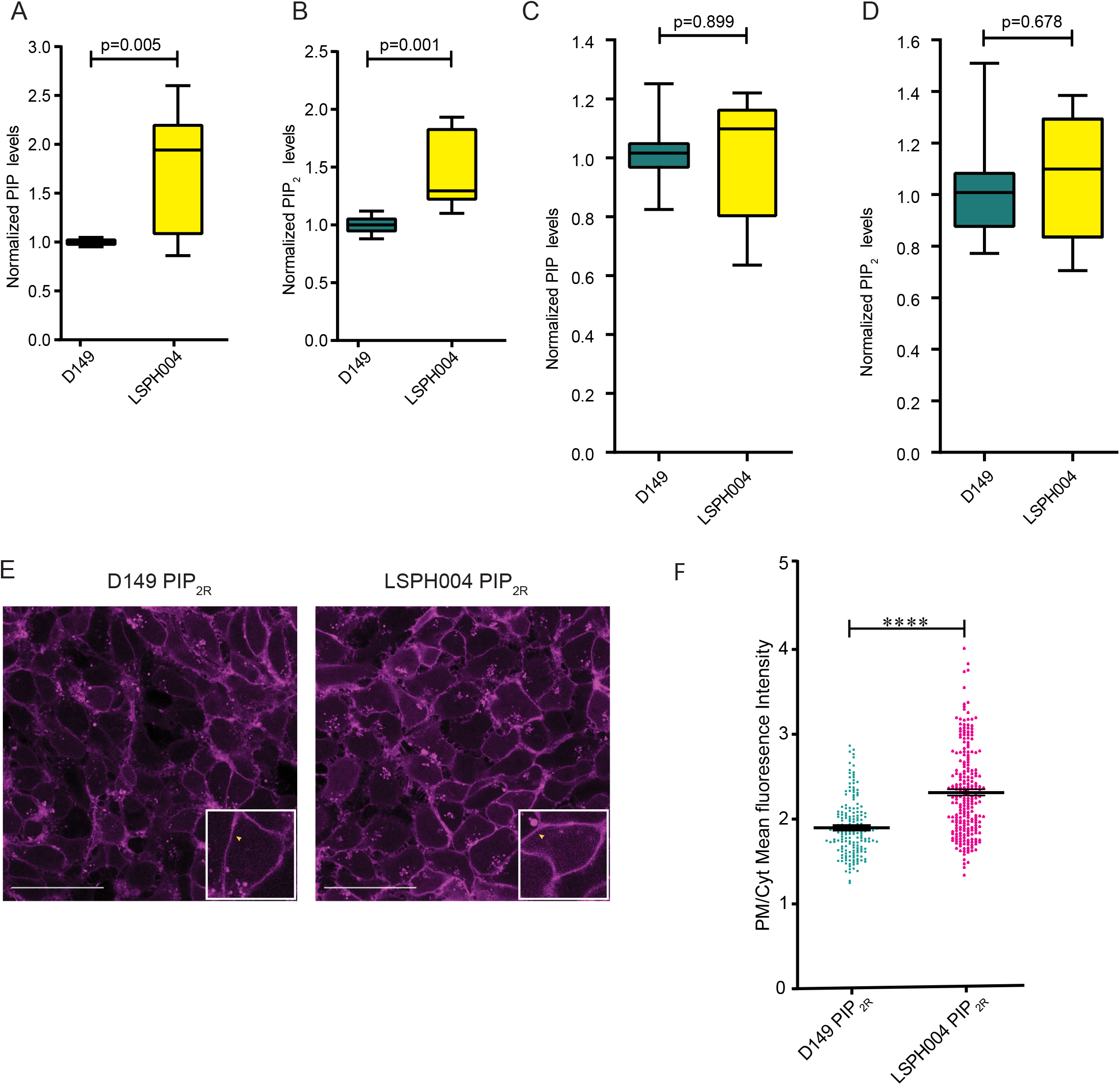
Mass spectrometry and biosensor estimation of PIP & PI(4,5)P_2_. (A) Box and Whisker Plot showing total PIP levels using LCMS in whole cell lipid extract from hiPSCs of Control (D149, n=9) and Patient lines (LSPH004, n=10). X-axis denoting samples and Y-axis represents normalized PIP levels. Statistical test: Two-tailed unpaired t-test with Welch’s correction. Whiskers at minimum and maximum values and a line at the median. (B) Box and Whisker Plot showing total PIP_2_ levels using LCMS in whole cell lipid extract from hiPSCs of Control (D149, n=9) and Patient lines (LSPH004, n=10). X-axis denoting samples and Y-axis represents normalized PIP_2_ levels. Statistical test: Two-tailed unpaired t-test with Welch’s correction. Whiskers at minimum and maximum values and a line at the median. (C) Box and Whisker Plot showing total PIP levels using LCMS in whole cell lipid extract from Neural Stem Cells of Control (D149, n=12) and Patient line (LSPH004, n=12). X-axis denoting samples and Y-axis represents normalized PIP levels. Statistical test: Two-tailed unpaired t-test with Welch’s correction. Whiskers at minimum and maximum values and a line at the median. (D) Total PIP_2_ levels using LCMS in whole cell lipid extract from Neural Stem Cells of Control (D149, n=12) and Patient line (LSPH004, n=12). X-axis denoting samples and Y-axis represents normalized PIP levels. Statistical test: Two-tailed unpaired t-test with Welch’s correction. Whiskers at minimum and maximum values and a line at the median. (E) Representative confocal maximum z-projections of mCherry::PH-PLCδ expressing PIP_2_ reporter lines used to estimate plasma membrane/cytosolic PIP_2_ probe fluorescence ratio (PM/Cyt). Enlarged insert shows mCherry::PH-PLCδ biosensor localization to the plasma membrane in a single cell. (F) Dot plot denoting quantification of PIP_2_ levels using mCherry::PH-PLCδ biosensor in Control (D149) and Patient (LSPH004) PIP_2_ reporter neural stem cell lines. X-axis denotes samples, Y-axis represents plasma membrane/cytosolic fluorescence ratio (PM/Cyt) of mCherry::PH-PLCδ biosensor. Each dot represents PM/Cyt obtained from a cell. Statistical test: Two-tailed unpaired t-test with Welch’s correction. **** p-value <0.0001. Error bars: S.E.M.

### Elevated plasma membrane PI(4,5)P_2_ levels in Lowe NSC

Since we did not observe a significant change in the total mass of PI(4,5)P_2_ in the LS patient NSC, we wondered whether the OCRL enzyme might control a relatively small but functional pool of PI(4,5)P_2_ at a specific endomembrane in NSC and changes in this small pool might not be reflected in measurements of total PI(4,5)P_2_ mass measurements. In cells, a key sub-cellular membrane where PI(4,5)P_2_ is enriched is the plasma membrane. To measure plasma membrane PI(4,5)P_2_ levels, we used the reporter lines expressing mCherry::PH-PLC-δ. Live cell imaging was performed on D149 and LSPH004 NSC expressing mCherry::PH-PLC-δ. We estimated PI(4,5)P_2_ levels at the plasma membrane by imaging the biosensor expressing cells in a monolayer and calculating the plasma membrane to cytosolic (PM/Cyt) fluorescence ratio of the probe (Fig. 4E). Quantification of these data revealed that the PI(4,5)P_2_ levels were significantly higher at the plasma membrane of the patient cell line LSPH004 compared to D149 (Fig. 4F). Thus, loss of OCRL alters the pool of PI(4,5)P_2_ at the plasma membrane in NSC.

### Compensatory mechanisms for loss of OCRL function

*OCRL* is part of a large family of lipid 5-phosphatases in the human genome (Ramos et al., 2019) and the function of this gene family in neurodevelopment has not been studied. Loss of OCRL function in LS patients might result in compensatory changes in the expression of other 5-phosphatase family members during brain development, thus leading to modest or no changes in PI4P and PI(4,5)P_2_ levels. To determine the expression pattern of these phosphatases during neural development, we performed qRT-PCR analysis for all the ten 5-phosphatases encoded in the human genome (Ramos et al., 2019) in hiPSC, NSC and 30 DIV neuronal cultures of D149. This analysis revealed an interesting and variable pattern of expression for each gene at these specific stages of neural differentiation *in vitro*. While some 5-phosphatases such as *SYNJ1, SYNJ2, INPP5B* and *INPPL1* were expressed at similar levels across all three stages, *INPP5D* appeared downregulated during neuronal differentiation. By contrast, a set of 5-phosphatases including *OCRL, INPP5F, INPP5K, INPP5E* and *INPP5J* all showed upregulation during neuronal differentiation (Fig. 5A). We then compared expression of all ten phosphatases in each of the three developmental stages hiPSC, NSC and 30 DIV neurons between D149 and LSPH004. At the hiPSC stage, five of ten 5-phosphatases that we assayed were upregulated in the patient line LSPH004; these were *INPP5D, INPP5E, INPP5F, INPP5K, INPPL1.* By contrast, *SYNJ1* and *INPP5J* were downregulated; as expected, *OCRL* was significantly down regulated in the patient hiPSC (Fig. 5B). In NSC, we observed five out of ten 5-phosphatases were upregulated in the patient line namely, *INPP5J, INPP5E, INPP5F, SYNJ2, INPPL1* and the only 5-phosphatase downregulated was *OCRL* (Fig. 5C). We also differentiated the NSC into neurons and compared expression levels of the phosphatases at 30 DIV between D149 and LSPH004; this revealed that except for a modest downregulation of *INPP5K*, there were no compensatory changes in LSPH004 (Fig. 5D). Thus, loss of *OCRL* results in distinctive patterns of compensatory changes in 5-phosphatase gene expression at various stages of neurodevelopment (Fig. 5E).

**Figure 5.**
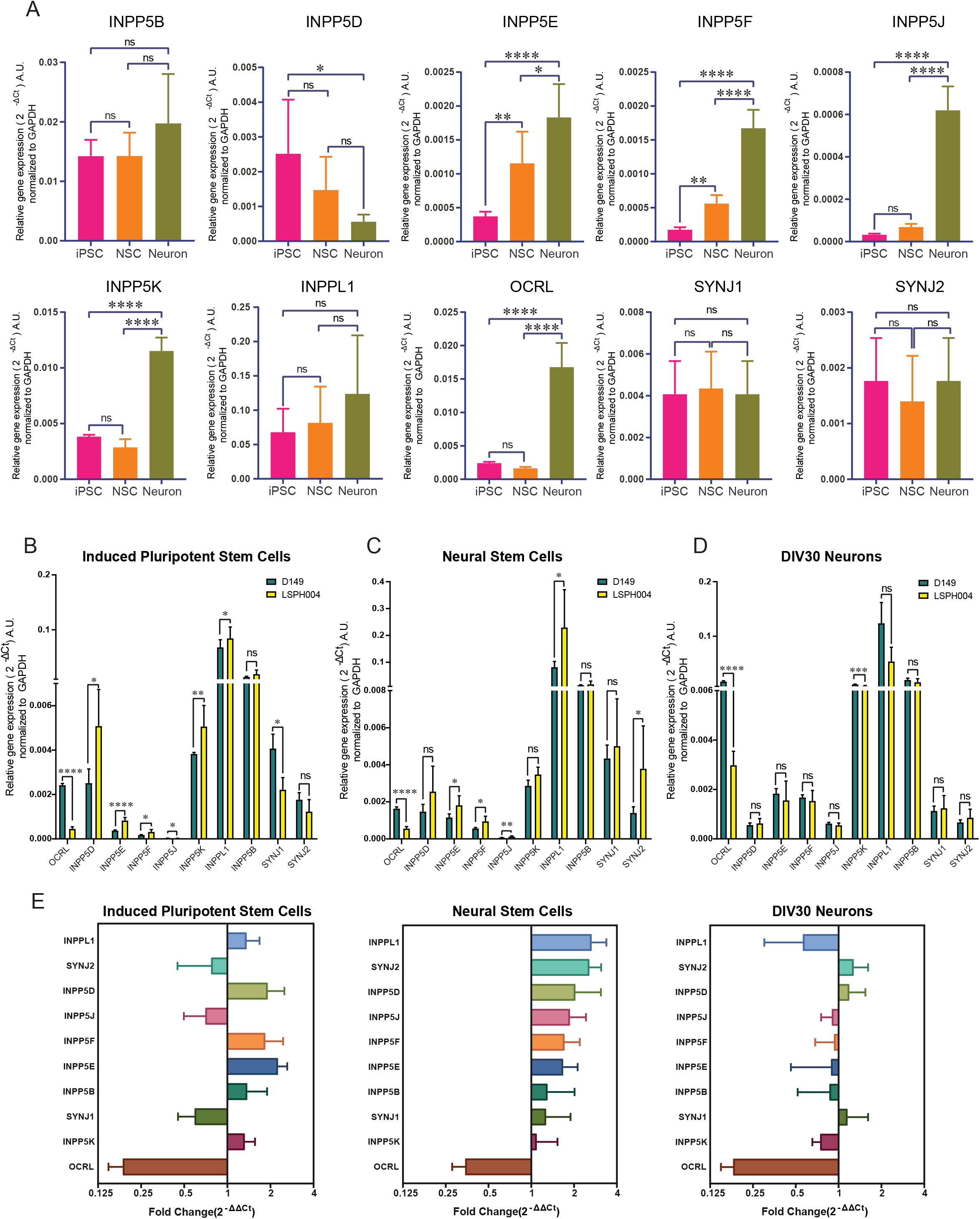
Expression analysis of inositol-5-phosphatase genes in hiPSC, NSC and 30 DIV neurons using qRT-PCR. (A) Quantitative real time PCR (qRT-PCR) showing mRNA expression of 10 inositol-5 - phosphatases in the human genome. Data is shown for hiPSC, NSC and 30 DIV neurons in the D149 control line. Statistical test: one-way ANOVA with post hoc Tukey’s multiple pairwise comparison. * p-value <0.05, ** p-value <0.01, **** p-value <0.0001. Error bars: Standard Deviation (B-D) Relative mRNA expression levels of 10 inositol-5 -phosphatases across hiPSC, NSC and 30 DIV neuron in D149 control line and LSPH004 patient line obtained from quantitative real time PCR (qRT-PCR). The expression levels have been normalized with GAPDH and the values represented in the terms of 2^-ΔCt^ on Y-axis. Statistical test: Two-tailed unpaired t-test with Welch’s correction. Error bars: Standard Deviation (E) Fold Change in expression of 10 inositol-5-phosphatases across hiPSC, NSC and 30 DIV neurons in LSPH004 patient line relative to D149 obtained from quantitative real time PCR (qRT-PCR) using the 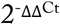method with GAPDH as a control gene. Error bars: Standard Deviation

### Transcriptomic changes in LS cells

To characterize the gene expression changes resulting from OCRL loss of function that may lead to neurodevelopmental defects, we performed transcriptomic analysis from hiPSC and NSC of D149 and LSPH004. At the iPSC stage, corresponding to the earliest stages of human embryonic development, we found ca. 475 genes upregulated and ca. 400 genes downregulated (Fig. 6A). Analysis of this set of altered genes using Gene Ontology revealed a number of categories of highly enriched downregulated genes suggestive of altered brain development (GO terms: generation of neurons, neurogenesis, neuron differentiation, nervous system development) (Fig. 6C-D, Supplementary Table 2). These included *SLITRK1, DCX, MAP2, BRINP1, CEP290, PAX7, NCAM1, SEMA6D* and *CNTN2*.

**Figure 6.**
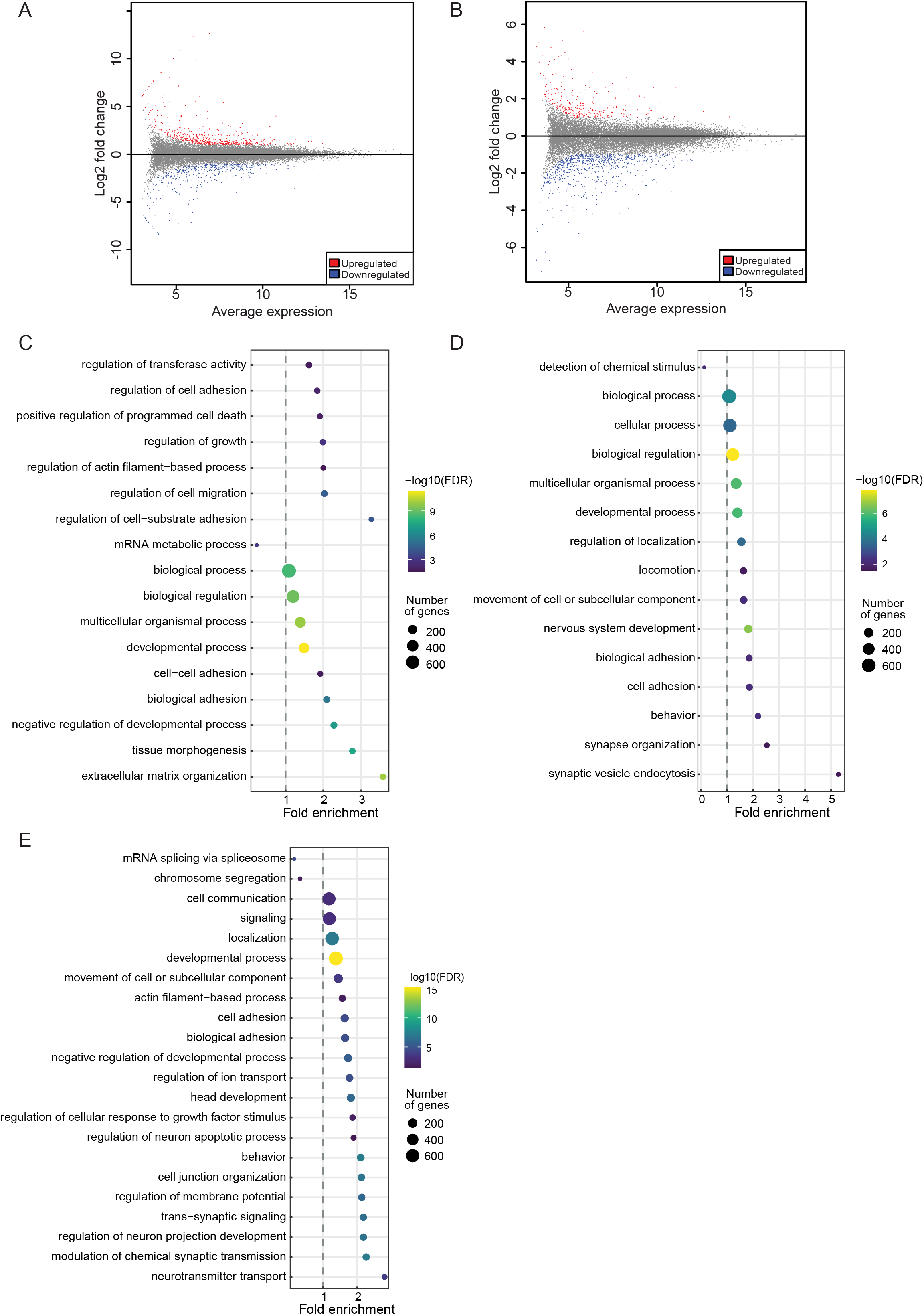
RNA-Seq analysis denoting DEGs and enriched Gene Ontology Biological Process terms in patient iPSCs and NSCs. (A) Scatter plot showing log_2_ fold change on Y-axis and mean of normalized expression counts of on X-axis from Differential Expression genes between control (D149) vs patient (LSPH004) hiPSC RNA-Seq. Red and blue dots indicate statistical significant up and down regulated genes respectively (-1.5 > Log_2_FC > 1.5, FDR ≤ 0.05) (B) Scatter plot showing log_2_ fold change on Y-axis and mean of normalized expression counts of on X-axis from Differential Expression genes between control (D149) vs patient (LSPH004) NSC RNA-Seq. Red and blue dots indicate statistical significant up and down regulated genes respectively (-1.5 > Log_2_FC > 1.5, FDR ≤ 0.05) (C–E) Dot-plot showing GO (Gene Ontology) Biological Process term enrichment, Y-axis shows enriched GO terms whereas X-axis shows Fold enrichment of each term in upregulated genes of hiPSC LSPH004 (C), downregulated genes of hiPSC LSPH004 (D) and downregulated genes of NSC LSPH004 (E). Colours indicate the –log_10_FDR from Fisher’s exact test, and dots size is proportional to the number of differentially expressed genes (DEG) in the given pathway.

Likewise, a comparative analysis of D149 and LSPH004 NSC revealed more than 750 downregulated and 250 upregulated genes (Fig. 6B). Gene Ontology analysis of this gene set also revealed strong enrichment of GO terms suggesting altered neural development with the most highly significant GO terms being neurogenesis and nervous system development and function (Fig. 6E, Supplementary Table 3). These include *SEMA3F, PLXND1, SOX8, LRRC7, HAPLN2, NTRK2, NRG1* and *SYT4.* These findings suggest a role for OCRL in the development of the brain, starting with the earliest cellular stages of embryogenesis, that can be analysed in this experimental system.

## Discussion

Although several sub-cellular functions have been described for OCRL (De Matteis et al., 2017; Mehta et al., 2014), the mechanism by which mutations in this gene result in human neurodevelopmental defects remains unknown. A particular challenge in understanding the brain phenotype in LS patients arises from the inability to obtain conventional biopsy samples from the human brain. This is in contrast to a limited number of studies where renal biopsies (De Leo et al., 2016) and skin samples (Wenk et al., 2003) have been used to address some specific aspects of the LS phenotype. In addition, since LS is neurodevelopmental in origin, understanding brain phenotypes present at birth requires the ability to study the cellular and physiological changes during brain development. To achieve this goal, it is necessary to have a model in which the development of human brain cells can be studied *in vitro*. A suitable model must recapitulate key aspects of brain development *in vitro* and also the key biochemical effects seen in LS patient cells.

We have identified a family with LS (Ahmed P et al., 2021) and in this study, report the generation of hiPSC from patients in this family as a resource to analyze the cellular and molecular basis of brain development in LS. Since *OCRL* encodes a PI(4,5)P_2_ phosphatase, it is expected that loss-of-function mutations in this gene should result in elevation of PI(4,5)P_2_ levels and a drop in PI4P levels. Our measurements of the mass of PIP_2_ and PIP in hiPSC cultures of the patient lines showed an elevation of total PIP_2_ and PIP in LSPH004 cells. This parallels observations on cultured human fibroblast cell lines from LS patients that have been previously reported (Wenk et al., 2003). In this study, we also engineered the LSPH004 line to express a fluorescent reporter for plasma membrane PI(4,5)P_2_ levels and found using this approach that the levels of this lipid at the plasma membrane of the patient line were elevated in comparison to control. Thus, the *in vitro* model system presented here recapitulates the key biochemical defect in phosphoinositide metabolism reported in LS patient tissues.

We differentiated these hiPSC lines using protocols based on the known principles of developmental neurobiology to generate 2D cultures of human neural tissue, primarily neurons *in vitro*. This developmental process *in vitro* recapitulates key aspects of brain development *in vivo* (Mertens et al., 2016). These hiPSC derived neural cultures show progressive increase of neural activity as a function of the age of the culture (DIV) (Fig. 2D, E). A comparison of neural activity development from LS patient derived cell lines reported here compared to those from control lines will provide an insight into the altered physiological development in brain tissue of LS patients. During metazoan development, gene expression is a key process that determines cell fate specification and differentiation. To understand gene expression changes that underlie the neurodevelopment phenotype in LS, we compared transcriptomes between control and LSPH004 at both the iPSC and NSC stages. Large differences in transcriptome were seen at both of these developmental stages (Fig. 6) revealing substantial changes in the expression of genes annotated as having roles related to nervous system development. These findings imply that the use of the hiPSC lines described here are likely to be valuable in understanding the mechanism by which loss of OCRL leads to altered brain development. Altogether, the model system described here allows an analysis of the physiological development in LS patients as well as the biochemical and molecular correlates of this process.

In addition to neuronal differentiation, the LS patient derived hiPSCs can be differentiated into glia to study the contribution of OCRL function in these non-neuronal cell types to the altered brain development in LS patients. Hyperechoic periventricular lesions have been described in MRI scans of patients with LS with features suggestive of enhanced gliosis (Sener, 2004) and evidence suggestive of enhanced gliosis in a fish model of LS has been reported (Ramirez et al., 2012). However, its relevance in the brain tissue of human LS patients remains unknown and the hiPSC lines generated here will also allow this question to be studied. In addition, the use of co-cultures of one neural cell type (e.g neurons) from LS patients with glial cells derived from controls and vice versa (G. Nadadhur et al., 2019) will allow an assessment of the cell autonomous and non-cell autonomous function of *OCRL* in supporting normal brain development. The developing brain is a complex 3D tissue and 2D cultures may not recapitulate all aspects of the cell-cell interactions or cortical layer formation that impact normal brain development. However, hiPSC can be used to generate 3D organoids of the developing brain (Di Lullo and Kriegstein, 2017); the use of the LS patient lines described in this study will also allow the role of OCRL in supporting the 3D architecture during brain development to be investigated. Finally, a recent study has proposed the use of the phosphoinositide 3-kinase inhibitor, alpelisib, for correcting renal defects in a mouse model of LS (Berquez et al., 2020). The availability of the hiPSC lines described in this study can help evaluate the effectiveness of such inhibitors in alleviating the phenotypes of LS in brain tissue.

It has been noted that although LS is a monogenic disorder, there is considerable variability in the brain phenotypes of individual patients. It has been proposed that such variability in clinical features between patients may arise from the impact of background mutations in the genomes of individual patients carrying functionally equivalent mutations in *OCRL*. However, the cellular and developmental basis of this has not been tested. The unique genetic structure and clinical features of the LS family we have studied (Ahmed P et al., 2021) provides a unique opportunity to address this scientific question. The three patients in this study, LSPH002, LSPH003 and LSPH004 vary clinically in their neurodevelopmental phenotype with LSPH004 showing much severe brain phenotype than LSPH002 and LSPH003 who are identical twins. In this study, we have generated hiPSC from all three patients; these cell lines carry the identical mutation in *OCRL* in the background of the genome of the individual patient from whom they were derived. By differentiating these hiPSC into neural tissue and comparing LSPH004 with LSPH002 and LSPH003, it is likely that we will discover the cellular and developmental correlates of the variable brain phenotype. Since the whole genome sequence of each line has been determined, it is also possible to experimentally test the importance of specific variants in the patient genome by CRISPR/Cas9 genome editing of specific variants (Paquet et al., 2016) followed by phenotypic analysis.

Although we observed changes in the levels of PIP and PIP_2_ in patient derived hiPSC from LS patient, several key aspects of altered phosphoinositide biochemistry in LS patient cells remain to be understood: (i) Changes were relatively modest and the changes in PIP and PIP_2_ mass seen in hiPSC were no longer evident at the NSC stage. These observations indicate a plasticity in the control of PI(4,5)P_2_ levels, perhaps by other enzymes that may also regulate PI(4,5)P_2_ levels that needs to be understood. (ii) In addition to elevated PIP_2_ levels in LS hiPSC, PIP levels were also elevated; this is unexpected though it has also previously been reported in fibroblasts from LS patients (Wenk et al., 2003). (iii) It is unclear if the elevated PIP_2_ levels, PIP levels or both lead to cellular effects leading to patient phenotypes. In order to address these questions, the system described here offers many advantages. Our genome engineering approach using the GSH in these lines offers the opportunity for controlled expression of enzymes to modulate the levels of phosphoinositides acutely (Idevall-Hagren and De Camilli, 2014; Varnai et al., 2006) to test specific hypotheses related to the role of individual lipids in altered cellular and developmental phenotypes in developing neural cells. One likely mechanism underlying plasticity in the control of PI(4,5)P_2_ levels in LS patient cells is changes in the expression of other genes encoding members of the 5-phosphatase family of enzymes. Indeed, RT-PCR analysis of the ten members of the 5-phosphatase in LSPH004 revealed that there were distinctive patterns of up or down regulation in the expression of the 5-phosphatase gene family members in wild type cells during development. Such patterns of 5-phosphatase expression and compensatory changes of these in LS patient cells during development may underlie the specific nature of the neurodevelopmental defects in LS patients. In summary, the hiPSC resources and their engineered derivatives described here offer powerful tools for understanding the regulation of PI(4,5)P_2_ to PI4P balance in the developing nervous system by OCRL and the mechanism by which loss of this activity leads to neurodevelopmental defects.

## Acknowledgements

This work was supported by the National Centre for Biological Sciences-TIFR, the Department of Biotechnology, Government of India through the Accelerator Program for Discovery in Brain Disorders (BT/PR17316/MED/31/326/2015), the Pratiksha Trust and a Wellcome-DBT India Alliance Senior Fellowship to PR (IA/S/14/2/501540). We thank the NCBS Imaging & Flow Cytometry, Genomics, High performance computing, Stem cell and Biosafety facilities for support. We thank Dr. O. Mukherjee and M. Rao for advice in stem cell generation.

## Materials and Methods

### Cell Lines and culture conditions

#### hiPSCs

D149 (Iyer et al., 2018), NIH5 (Baghbaderani et al., 2015), LSPH002, LSPH003, LSPH004 (generated in this study). When grown on a mouse embryonic fibroblast (MEF) feeder layer, all hiPSC were grown in standard HuES (Human embryonic Stem cell) media. When transitioned to feeder-free extra cellular matrix Matrigel (hESC-qualified Matrigel Corning, #354277) coated surface, the cells were grown in E8 complete media (E8 basal + E8 supplement). Cultures were maintained at 37°C and 5% CO_2_ throughout.

#### NSCs

D149, LSPH004, D149-GSH, LSPH004-GSH, D149-PIP_2R_, LSPH004-PIP_2R_ (generated in this study). All Neural Stem Cells (NSCs) were grown on Matrigel coated tissue culture plasticwares in neural expansion media at 37°C and 5% CO_2_.

### Mycoplasma testing

Mycoplasma contamination was checked using spent media from sub-confluent hiPSC and NSC dishes after 48 h in culture. MycoAlert™ (Lonza, #LT07–418) was used per the manufacturer’s protocol.

### Karyotyping

Overall chromosomal integrity of hiPSCs and NSCs was confirmed by karyotyping. For metaphase preparation, cells were arrested in log phase by treating with 0.1 µg/mL Colcemid™ (Gibco, #15212- 012) treatment for 45 min at 37°C. Cells were harvested in fresh Carnoy’s fixative (Methanol:Glacial Acetic Acid at 3:1) and G-banding karyotype analysis performed at a National Accreditation Board for Testing and Calibration Laboratories, India (**NABL**) accredited facility.

### Generation of hiPSC lines

Blood was drawn from donors after informed consent and under aseptic conditions following IRB regulations. A peripheral blood mononuclear cell fraction was obtained by density gradient centrifugation and transformed into Lymphoblastoid Cell Lines (LCL) by Epstein Barr virus (EBV) transformation using previously established protocol (Hui-Yuen et al., 2011). These LCLs were reprogrammed by electroporation of plasmids containing Yamanaka factors to generate hiPSCs as previously described (Iyer et al., 2018). On-feeder hiPSC cultures were gradually transitioned to feeder-free conditions by weaning off from the standard HuES media to Essential 8™ medium (Gibco, #A1517001) and maintained on hESC-qualified Matrigel coated surface. hiPSCs were frozen at a density of 1×10^6^ in 500µL PSC cryomix (Gibco, #A26444-01). The plasmid footprint in feeder-free iPSC lines LSPH002 (passage 7), LSPH003 (passage 7) and LSPH004 (passage 8) was tested using PCR analysis with relevant primer sets (Supplementary Table 5), with 3% DMSO as a PCR additive. Plasmid pCXLE-hUL was used as a positive control while iPSC NIH5 was used as negative control. To ascertain differentiation potential, embryoid bodies (EB) were generated by transferring hiPSCs to a non-adherent surface in standard HuES media without bFGF for 48 h, and cDNA probed for primers specific for the three lineages.

### Generation of NSCs

NSCs were generated as previously described (Mukherjee et al., 2019) with slight modifications. Briefly, hiPSCs were differentiated to form embryoid bodies in E6 medium (Gibco, #A1516401). Primary neural rosettes formed by this method were selected and manually passaged to obtain secondary and tertiary rosettes that were eventually triturated and plated as NSC monolayer in Neural Expansion Medium (NEM). As a measure to eliminate any non-NSC cells and obtain a reliable homogeneous NSC culture, the generated NSCs were subjected to CD133+ selection as previously described via fluorescence-activated cell sorting (FACS) (Peh et al., 2009). A sub-confluent NSC culture maintained in NEM was enzymatically dissociated using Stempro Accutase (Gibco, #A11105-01) and following a wash in PBS, cells were immunolabelled using CD133-PE conjugated primary antibody (Abcam, #ab253271) at 10μl antibody/million cells and incubated at room temperature for 30 min in dark. Cells were washed with PBS and resuspended in 1mL sorting media [(1x DMEM/F12-without phenol red (Gibco, #21041025), 1% FBS (Gibco, #16000-044) + Penicillin-Streptomycin (Gibco, #15140-122)] keeping the concentration at 2 million cells per mL to obtain efficient sorting. The cells were sorted using FACS- Aria-III instrument (BD Biosciences) (Supplementary Fig.1D-E). Forward and side scatter parameters were adjusted so as to eliminate cell clumps and debris. Cells with highest fluorescence intensity were collected and plated on Matrigel at a concentration of 0.5 million cells per well of 12-well tissue culture plate in NEM and expanded. Cells were periodically checked for bacterial or mycoplasma contamination. NSCs were further characterized by immunofluorescence and neuronal differentiation.

### Reporter line generation

For generation of D149 and LSPH004 NSC safe harbor lines, cells growing in Neural Expansion Medium (NEM) without Penicillin-Streptomycin were harvested by enzymatic dissociation using Stempro Accutase for 5 min at 37 °C. The cells were spun down at 1000g for 3 min to remove Accutase post neutralization at room temperature. After a wash with PBS, cells were incubated in buffer-R (included in Invitrogen, #MPK1025) with 2 μg/μL each of plasmids: AAVS-1-TALEN-R (ZYP017), AAVS-1-TALEN-L (ZYP018) and ZYP037 (Pei et al., 2015) for 3 min at a concentration of 1 million cells per hit. The cell-plasmid mixture was electroplated using the Neon Electroporation System (Invitrogen, USA. #MPK1025) as per manufacturer’s protocol under the following standardized parameters (1100 volts, width-20, pulses-2) and plated in 1 well of 12well plate coated with Matrigel containing pre warmed NEM without Penicillin-Streptomycin. A complete media change was performed 4 h post electroporation to remove dead cells and debris which could potentially cause cytotoxicity in the culture if not removed. After 24 h, cells were checked under an epifluorescence microscope for expression of GFP and upon confirmation, the cells were subjected to Puromycin (Gibco, #A1113803) selection at a concentration of 0.4 μg/mL for 7-9 days with daily media change. After a week, large GFP positive colonies were visible in the plate, at this point the culture was subjected to Fluorescence-activated cell sorting (FACS) to remove any GFP-negative cell. The mixed cell culture was enzymatically dissociated using Stempro Accutase, washed once with PBS and resuspended in sorting media [(1x DMEM/F12-without phenol red (Gibco, #21041025), 1% FBS + Penicillin-Streptomycin] at a concentration of 2 million cells per mL. The cells were sorted using FACS- Aria- III (BD Biosciences). Forward and side scatter parameters were adjusted so as to eliminate cell clumps and debris. The gating parameter threshold for GFP-positive cells was set using non-electroporated control NSCs. The cells were collected in NEM and plated in pre-incubated Matrigel coated plate at a concentration of 0.5 million cells per well of 12-well tissue culture plate in NEM and expanded for cryopreservation.

The Safe Harbor NSCs; D149-GSH and LSPH004-GSH were enzymatically dissociated using Stempro Accutase for 5 min at 37 °C. The cells were spun down at 1000g for 3 min at room temperature to remove Accutase. After a wash with PBS they were resuspended in Buffer-R (included in Invitrogen, #MPK1025) with 2 μg/μL each of plasmids: LoxP-CAG-mCherry::PH-PLCδ (ZYP070-PIP_2R_) and Cre-Recombinase (ZYP073) at a concentration of 1 million cells per hit of electroporation which was performed using Neon Electroporation System (Invitrogen, USA. #MPK1025) as per manufacturer’s protocol under the following standardized parameters (1100 volts, width-20, pulses-2) and plated in 1 well of 12-well plate coated with Matrigel containing pre warmed NEM without Penicillin-Streptomycin. A complete media change was performed 4 h post electroporation to remove dead cells and debris which could potentially cause cytotoxicity in the culture if not removed. After 24 h, cells were checked under epifluorescence microscope for expression of mCherry and upon confirmation, the cells were subjected to G418 (Gibco, #10131035) selection at a concentration of 400 μg/mL for 15-20 days with daily media change. After about two weeks under selection, distinct mCherry positive colonies were observed, at this point the culture was subjected to Fluorescence-activated cell sorting (FACS) to remove GFP-positive cells. The cells were sorted at a concentration of 2 million cells per mL using FACS- Aria-III (BD Biosciences). Forward and side scatter parameters were adjusted so as to eliminate cell clumps and debris. The gating parameter threshold for mCherry-positive cells was set using GFP-positive and non-electroporated control NSCs. Post FACS cells were collected in NEM and plated in pre-incubated Matrigel coated plate at a concentration of 0.5 million cells per well of 12-well tissue culture plate in NEM and expanded to freeze down additional stock vials.

### qRT-PCR

#### Isolation of RNA and cDNA synthesis

Total RNA was extracted from well characterized hiPSCs and NSCs of patient and control lines using TRIzol (Ambion, Life Technologies, #15596018) as per manufacturer’s protocol in 6 biological replicates and quantified using a Nanodrop 1000 spectrophotometer (Thermo Fisher Scientific). Following treatment with 1U of DNase I (amplification grade, Thermo Fisher Scientific, #18068-015), 1 μg of the RNA from each replicate was used for cDNA synthesis in a reaction mixture containing 10 mM DTT and 40U of RNase inhibitor (RNaseOUT, Thermo Fisher Scientific, #10777-019). The reaction mixture of 45.5 μl was incubated at 37°C for 30 min followed by heat inactivation at 70°C for 10 min, following which 200U of Superscript II Reverse Transcriptase (Invitrogen, #18064-014) was added to the reaction volume along with 2.5 μM of random hexamers, and 0.5 mM of dNTPs making the final volume to 50 μl. The reaction was then incubated at 25°C for 10 min, followed by 42°C for 60 min and then heat inactivated at 70°C for 10 min on ProFlex PCR Systems (Life Technologies).

#### Real-Time Quantitative PCR

The primers used for qRT-PCR were designed using Primer-BLAST, NCBI (https://www.ncbi.nlm.nih.gov/tools/primer-blast/). With a condition of spanning exon-exon junction of their respective genes, their details are provided in supplementary Table 5. Real-Time qRT-PCR was performed in a volume of 10μl with Power SYBR Green Master mix (Applied Biosystems, #4367659) on an Applied Biosystems ViiA7 system. It was performed with technical triplicates from the patient and control lines with primers for genes of interest and GAPDH was used as a house- keeping gene control. A no-reverse transcriptase control was also set up. The reaction was run under the following conditions: 50°C for 2 min, 95°C for 10 min, followed by 40 cycles of 95°C for 30 sec (denaturation), 60°C for 30 sec (annealing) and 72°C for 45 sec (extension). The C_t_ values obtained for individual genes were normalized to those of GAPDH from the same sample. The relative expression levels were calculated using ΔC_t_ method, whereas the fold change between patient and control was calculated using ΔΔC_t_ method.

#### Junction PCR

Junction PCR was performed using the genomic DNA of the edited cells, extracted using QIAamp DNA Mini Kit (Qiagen, #51304) using manufacturer’s protocol and quantified using a Nanodrop 1000 spectrophotometer (Thermo Fisher Scientific). Insertion of the ZYP037 donor template was confirmed using two pairs of primers; one of the primers in each pair annealed outside the region spanned by the homology arm, in the AAVS1 locus, while the other annealed within the inserted template, to avoid false detection of residual plasmid if any. 50 ng of genomic DNA was used as the template in 20 μl PCR reaction consisting Phusion Pol (0.02 U/µl), 25 mM DNTPs, 5X HF-buffer and appropriate primers. The reaction was run as a touchdown PCR in two steps as follows: 98°C for 3 min (initial denaturation), step-1 [98°C for 20 sec-denaturation, 66°C to 61°C (-0.5°C per cycle) for 30 sec- annealing, 72°C for 2 min-extension] ×10 cycles, step-2 [98°C for 20 sec-denaturation, 55°C for 30 sec- annealing, 72°C for 2 min-extension] x25 cycles, 72°C for 5 min (final extension).

### List of Primers and Antibodies used in this study is provided as Supplementary Table 5

#### Immunocytochemistry

hiPSCs were characterized for pluripotency markers SOX2 and SSEA4 using the PSC immunocytochemistry kit (Invitrogen, #A24881) as per manufacturer’s instructions.

NSC cultures were fixed using 4% formaldehyde in phosphate-buffered saline (PBS) for 20 min, permeabilized using 0.1% Tx-100 for 5 min and incubated at room temperature for 1 h in a blocking solution of 5% BSA in PBS. Primary antibodies at respective dilutions were added and incubated overnight at 4°C in blocking solution, followed by incubation with secondary antibodies in blocking solution (Invitrogen) for 1 h.

Confocal images were recorded by collecting a range of z-stack using an Olympus FV 3000 confocal microscope. Epifluorescence images were captured using EVOS® FL Cell Imaging System (Thermo Fisher Scientific). The image stack was merged using Z-project (maximum intensity projection) function using ImageJ (National Institute of Health, USA, http://imagej.nih.gov/ij).

### Quantitative FACS analysis

hiPSCs were characterized quantitatively for pluripotency markers OCT4 and SSEA4 by flow cytometry. Briefly, the cells were detached using Stempro Accutase for 5 min at 37°C and washed with standard HuES media. The cells were then washed once with PBS (Phosphate-buffered saline) and fixed with 4% PFA solution at a cell density of 1 × 10^6^ cells per mL at room temperature for 15 min. The cells were then permeabilized with 0.1% v/v Triton X-100 in HBSS for 15 min at room temperature, washed once with HBSS and incubated with 1% BSA in HBSS for 30 min. Following this, the cells were incubated with primary antibodies in 1% BSA in combination at the following dilutions, Rabbit Anti-OCT4 (Thermo Fisher Scientific, #A24867) at 1:100 and Mouse IgG3 Anti- SSEA4 (Thermo Fisher Scientific, #A24866) at 1:100 and incubated for 3 h at room temperature with gentle mixing. The cells were then washed 2-3 times in HBSS and incubated with appropriate secondary antibodies, Alexa Fluor™ 594 donkey anti-rabbit Antibody (Thermo Fisher Scientific, #A24870) and Alexa Fluor™ 488 goat anti-mouse IgG3 (Thermo Fisher Scientific, #A24877) for 1 h at room temperature. Before use, the secondary antibodies were diluted at 1:250 in 1% BSA. The cells were washed twice and then analyzed using a BD FACSVerse cytometer. (Supplementary Fig. 1A-C)

### Western Blotting

NSCs and 30 DIV neurons were harvested using Stempro Accutase and pelleted at 1000g for 5 min then washed thrice with ice-cold PBS. The pelleted cells were homogenized in 1X RIPA lysis buffer containing freshly added phosphatase and protease inhibitor cocktail (Roche). To remove cellular debris, crude RIPA lysates were centrifuged at 13,000 rpm for 20 min at 4 ° C. The supernatant was transferred to a new tube and quantified with a Pierce BCA protein assay (Thermoscientific, #23225). Thereafter, the samples were heated at 95°C with Laemmli loading buffer for 5 min and 20 ug protein was loaded onto Bolt™ 4 to 12%, Bis-Tris SDS gel (Invitrogen, #NW04120BOX). The proteins were then transferred onto a nitrocellulose membrane and incubated overnight at 4°C with indicated antibodies. The blots were then washed thrice with Tris Buffer Saline containing 0.1% Tween-20 (0.1% TBS-T) and incubated with 1:10,000 concentration of appropriate HRP-conjugated secondary antibodies (Jackson Laboratories, Inc.) for 45 min. After three washes with 0.1% TBS-T, blots were developed using Clarity Western ECL substrate (Biorad) on a GE ImageQuant LAS 4000 system.

### Calcium imaging

Calcium imaging was performed in 30 DIV D149 and LSPH004 neurons, according to our previously published protocol with minor modifications (Sharma et al., 2020). Briefly, neurons were washed with Tyrode’s buffer solution (5 mM KCl, 129 mM NaCl, 2 mM CaCl_2_, 1 mM MgCl_2_, 30 mM glucose and 25 mM HEPES, pH 7.4) for 10mins and later incubated with 4 uM fluo-4/AM (1 mM, Molecular probes, #F14201) and 0.002% pluronic F-127 (Sigma, #P2443) in the Tyrode’s buffer solution in dark for 30-45 min at room temperature. Following dye loading, the cells were washed again with the buffer thrice, each wash for 5 min. Finally, cells were incubated for an additional 20 min at room temperature to facilitate de-esterification. Ca^2+^ imaging was performed for 8 min with a time interval of 1 second using CellSens Dimension software (Olympus, build 16686) at 20X objective of wide-field fluorescence microscope Olympus IX-83. A four-minute baseline measurement was recorded to visualize calcium transients, followed by the addition of TTX to abolish calcium transients for another 4 min. Calcium traces were obtained using the CellSens Dimensions software by drawing a region of interest (ROI) manually around each neuronal soma. To plot the calcium traces, the raw fluorescence intensity values from each neuron were normalized to the first fluorescence intensity signal of the baseline recording. GraphPad Prism 5.0 was used to plot calcium traces.

### Electrophysiology

Whole-cell patch clamp recordings were performed at 10, 20, 30 and 40 DIV following initiation of neuronal differentiation of NSC. Recording micropipettes (5-7MΩ) were filled with internal solution composed of (in mM): 130 K-gluconate, 0.1 EGTA, 1 MgCl_2_, 2 MgATP, 0.3 NaGTP, 10 HEPES, 5 NaCl, 11 KCl, and Na^2^- phosphocreatinine (pH 7.4). Recordings were made at room temperature using a Multiclamp 700B amplifier (Molecular devices, USA). Signals were amplified and filtered at 10 Hz and 3 Hz, respectively. Voltage was corrected for liquid junction potential (−14 mV). The bath was continuously perfused with oxygenated artificial cerebrospinal fluid (ACSF) composed of (in mM): 110 NaCl, 2.5 KCl, 2 CaCl_2_, 10 glucose and 1 NaH_2_PO_4_, 25 NaHCO_3_, and 2 MgCl_2_ (pH 7.4) (Gunhanlar et al., 2018). For current-clamp recordings, voltage responses were evoked from a holding potential of −60mV to -70mV using 500 ms steps ranging from −5 to +70 pA in 5 pA intervals. Action potential (AP) properties were calculated from the first evoked AP in response to a depolarizing step. Repetitively firing neurons were defined as those capable of firing **>3 APs** in response to a depolarizing current step.

### Mass spectrometry

#### Lipid standards

17:0-20:4 PI(4,5)P2 (Avanti Lipids – LM1904), 17:0-20:4 PI(4)P (Avanti Lipids – LM1901), 17:0- 14:1 PE (Avanti Lipids – LM 1104)

### Solvent mixtures

**Quench mixture –**MeOH/CHCl_3_/1M HCl in the ratio 484/242/23.55 (vol/vol/vol).

**Lower Phase Wash Solution (LPWS) –**MeOH/1M HCl/CHCl_3_ in the ratio 235/245/15 (vol/vol/vol).

**Post derivatization Wash Solution (PDWS) –** CHCl_3_/MeOH/H2O in the ratio 24/12/9 (vol/vol/vol). Shake the mixture vigorously and allow settling into separate phases and use the upper phase only for washes.

### Lipid Extraction

Cells in culture were washed with DMEM/F12 to remove debris and then harvested using Stempro Accutase over a few minutes at 37°C. The Accutase was then neutralized and the cell suspension was then centrifuged at 1000g (for iPSCs) or 2500g (for NSCs) to pellet down the cells. The supernatant was discarded and the cell pellet was resuspended in 1 mL of 1X Phosphate Buffer saline (PBS) for a wash and transferred to a 2 mL low-bind tube. The tubes were centrifuged at the previously indicated speeds to pellet the cells again. The supernatant was discarded and the cell pellet was resuspended in 340 µl of 1X PBS and divided into two aliquots of 170 µl each for subsequent processing.

To each tube, 750 µl of ice-cold quench mixture, followed by 15 µl of a pre-mixed ISD mixture containing 25 ng of 37:4 PIP, 25 ng of 37:4 PIP_2_ and 50 ng of 31:1 PE was added. The tubes were vortexed for 2 min at about 1500 rpm. Thereafter, 725 µl of CHCl_3_ and 170 µl of 2.4 M HCl was added. The tubes were again vortexed for 2 min and kept at room temperature on the bench for 5 min. Two separate phases can be seen with a whitish precipitate at this stage.

All tubes were then spun at 1500 rpm in a benchtop centrifuge for 3 min to clearly separate the phases. In fresh 2 mL low-bind tubes, one for each sample, 708 µl of LPWS was added and kept aside. Once the phases were observed to be separate in the original tubes upon centrifugation, 900 µl of the lower organic phase was pipetted out by piercing through the upper phase and the protein precipitate at the interface and added to the tubes containing the LPWS. The tubes were then vortexed for 2 min and spun at 1500 rpm in a benchtop centrifuge for 3 min to separate the phases. The lower phase was pipetted out to extent possible taking care not to aspirate any of the upper phase and collected into a fresh tube.

### Lipid Derivatization

The following steps were performed inside a chemical hood, while wearing appropriate respirator mask. 50 µl of TMS-Diazomethane was added to each sample and incubated on a shaker at 600 rpm for 10 min at room temperature. At the end of 10 min, TMS-Diazomethane in each tube was quenched using 10 µl of Glacial Acetic acid. The tubes were inverted a few times to complete the quenching and carefully snapped open once to let the Nitrogen released during quenching to escape. At this point, the samples were moved out of the hood.

500 µl of the upper phase of PDWS was added to each sample. Tubes were then vortexed for 2 min and spun at 1500 rpm in a benchtop centrifuge for 3 min to separate the phases. 400 µl of the upper phase was discarded and another 500 µl of upper phase of PDWS was added to each tube and the vortex and spin steps were repeated as done earlier to separate the phases. Finally, the entire upper phase was discarded from each sample. 45 µl MeOH and 5 µl H2O was added to each tube, mixed and spun down. All the samples were then dried in a SpeedVac at 500 rpm for around 2 h till only about 10-20 µl of solvent was remaining. 90 µl of MeOH was added to reconstitute the sample to a final volume of about 100-110 µl and taken for injection and analysis.

### Liquid Chromatography and Mass Spectrometry

Samples were injected in duplicates. We performed chromatographic separation on an Acquity UPLC BEH300 C4 column (100 X 1.0 mm; 1.7 µm particle size - Waters Corporation, USA) using a Waters Aquity UPLC system connected to an ABSCIEX 6500 QTRAP mass spectrometer for ion detection. The flow rate was set to 100 µl /min.

Solvent gradients were set as follows –

Solvent A (Water + 0.1% Formic Acid)

Solvent B (Acetonitrile + 0.1% Formic acid)

0-5 min: 55% Solvent A + 45% Solvent B

5-10 min: Solvent B increased from 45% to 100%,

10-15 min: Solvent B at 100%,

15-16 min: Solvent B reduced from 100% to 45%,

16-20 min: 55% Solvent A + 45% Solvent B.

On the mass spectrometer, we first employed Neutral Loss Scans during pilot standardization experiments on biological samples and searched for parent ions that would lose neutral fragments corresponding to 490 a.m.u and 382 a.m.u indicative of PIP_2_ and PIP species respectively and likewise 155 a.m.u for PE species as described in (Clark et al., 2011). Thereafter, we quantified PIP, PIP_2_ and PE species in biological samples using the selective Multiple Reaction Monitoring (MRM) method in the positive ion mode (Supplementary Table 4). For each sample, PE levels measured were used to normalize for total cellular phospholipid content on the assumption that PE levels are not likely to be different between control and Lowe syndrome cell lines based on previous studies. The Sciex MultiQuant software was used to quantify the area under the peaks. For each run, the area under curve for each species of PIP, PIP_2_ and PE was normalized to PIP, PIP_2_ and PE internal standard peak respectively to account for differences in loading and ion response. Thereafter, the sum of normalized areas for all the species of PIP or PIP_2_ was then divided by the sum of normalized areas for all the species of PE in each of the biological samples to account for differences in total phospholipid extracted across samples.

### Live cell imaging of PI(4,5)P_2_ probe

5-10,000 NSC of LSPH004- PIP_2R_ and the control line D149- PIP_2R_ were seeded on 35mm glass bottom 15mm bore confocal dishes (Biostar LifeTech LLP #BDD011035) in NEM and cultured for 48 h to reach a uniform confluency. Prior to imaging, NEM was aspirated out of the dishes and cells were incubated with Hoechst (Invitrogen, USA. #H3570) at 5 μM final concentration in NEM for 10 min to stain the nucleus. The cells were then washed with PBS and NEM was added prior to imaging. Confocal images were recorded by collecting a z-stack using an Olympus FV 3000 confocal microscope. This was performed in 3 independent biological replicates.

### Image Analysis

The quantification of mCherry::PH-PLCδ probe was performed manually by generating the maximum z-projections of middle few planes of cells from confocal slices. Thereafter, line profiles were drawn across clearly identifiable plasma membrane regions and their adjacent cytosolic regions and the ratios of mean intensities (PM/Cyt) for these line profiles were calculated for each cell and used to generate statistics (Sharma et al., 2019).

### Transcriptomics

#### NGS sequencing

Total RNA was extracted from well characterized hiPSCs and NSCs of patient and control lines using TRIzol (Ambion, Life Technologies, #5596018) as per manufacturer’s protocol in 2 biological replicates. The RNA was quantified using Qubit4 dsDNA HS Assay Kit (Thermo Fisher Scientific, #Q32854) and run on a Bio-analyzer chip (Agilent High Sensitivity DNA Chip, #5067-4626) to assess integrity. Post NEB Next Poly(A) mRNA Magnetic Isolation Module (#E74906), 150 ng of total RNA (RIN values > 9) was used per sample for the library preparation using the NEB Next Ultra II Directional RNA Library Prep Kit for Illumina (New England Biolabs, #E7760L). The libraries were then sequenced on Illumina HiSeq 2500 sequencing platform using 2×100 bp sequencing format.

### Bioinformatics Analysis

Illumina sequenced paired-end reads were obtained from sequencing as mentioned above. The quality of processed reads (adapter removal and trimming) were evaluated using FastQC (version: 0.11.9). The RNA-seq reads were then mapped onto to the human reference genome (hg38) using HISAT2 (version: 2.1.0), the resulting BAM files containing the aligned reads were provided to HTSeq (version: 0.12.4) to obtain gene-level read count table using the reference annotation file (GTF format). We further utilized the iDEP (0.91)(Ge et al., 2018) to transform the read counts data using the regularized log (rlog) transformation method that is originally implemented in the DESeq2 [https://bioconductor.org/packages/release/bioc/html/DESeq2.html]. Differential expression, Enrichment analysis and Gene Ontology (Biological Process) were then conducted using DESeq2 and default settings within the iDEP (0.91). Genes with log_2_FC greater than +1.5 and lesser than -1.5 were considered as up and down regulated respectively while adhering to p-value < 0.05. Such filtered genes were used for further downstream analysis which included gathering enriched Gene Ontology (Biological Process) terms, for which the filtered gene list was provided to ShinyGO v0.66 (Ge et al., 2020) (http://bioinformatics.sdstate.edu/go/) with default filtering parameters. To remove redundant GO terms, we evaluated the gene lists for statistical significance by Fisher’s test using PANTHER (Mi et al., 2021) (http://pantherdb.org/). With a FDR <0.05 cut-off, we obtained the enrichment list which was further submitted to REViGO (Supek et al., 2011) (http://revigo.irb.hr/) which refines the number of redundant functional terms based on semantic similarity between the ontology terms. Using a tight filter of dispensability < 0.05, the Gene Ontology (Biological Process) terms were obtained.

### DNA constructs

ZYP017- AAVS1-TALEN-R (XCell Science. Novato, CA); ZYP018- AAVS1- TALEN-L (XCell Science. Novato, CA); ZYP037- AAVS1P-CAG-copCFP iRMCE (XCell Science. Novato, CA); ZYP070- DCX-GFP donor (XCell Science. Novato, CA); ZYP073- AAV-PGK-Cre (XCell Science. Novato, CA). To drive the probe expression in neural stem cells we replaced neuron specific DCX promoter in ZYP070 with CAG promoter by using the SalI and BsrGI restriction sites. mCherry::PH- PLCδ was cloned into ZYP070 at the GFP site by overlapping primers using GIBSON assembly.

### Statistics and Software

Two-tailed unpaired student’s t-test was used to compare datasets of two. One-way ANOVA with post hoc Tukey’s multiple pairwise comparison was used whenever the experiment consisted of more than two biological groups. All statistical analyses were performed on Graph Pad Prism (version. 8). Schematics were created with biorender.com.

## Data availability

RNA sequencing data used in this study has been submitted to the Sequence Read Archive database on NCBI under BioProject ID PRJNA741484

## Ethics Approval

This work was carried out under the ethics approval provided by the Institutional Ethics Committee, St. John’s Medical College & Hospital, Bangalore (IEC Study Ref. No. 28 / 2017) and the Institutional Ethics Committee, National Centre for Biological Sciences, Bangalore (NCBS/IEC-8/002). Institutional Stem Cell Committee Approval-National Centre for Biological Sciences (01/ICSCR/IX- 06.0l.2020-RP2).

## Author contributions

Conceptualization: PR, AV

Methodology: PR, AV

Software: -

Validation: BA, PB, SA, SS, YS, KG, ABS, AV

Formal analysis:BA,SS

Investigation: BA, PB, SA, SS, YS, KG, ABS, AV

Resources: AV

Data curation: BA, PB, SA, SS, YS, KG, ABS, AV

Writing - original draft: BA, PR

Supervision: PR

Project administration: PR

Funding acquisition: PR

